## Supplementary information for "The TOR signaling pathway regulates vegetative development, aflatoxin biosynthesis, and pathogenicity in *Aspergillus flavus*"

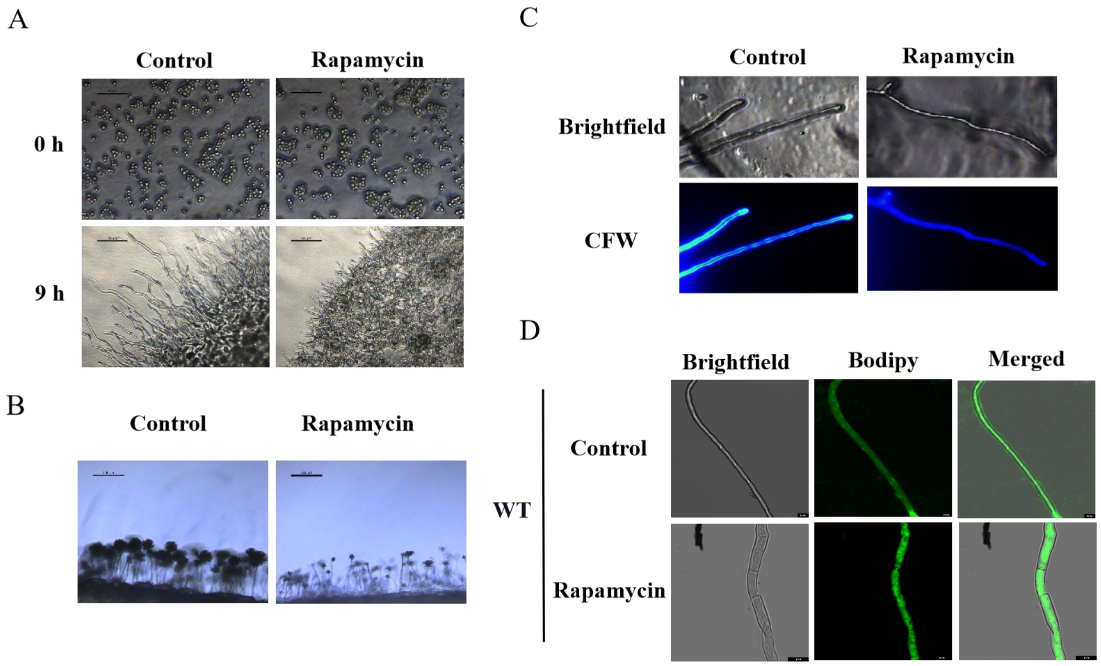


**Figure S1. Impacts of rapamycin on spore germination, conidiophore formation, and lipid droplet biogenesis of *A. flavus*.**

(A) Microscopic view of the spore germination of the WT strain treated with 100 ng/mL rapamycin.

(B) Microscopic view of the conidiophore formation of the WT strain treated with 100 ng/mL rapamycin.

(C) The hyphae treated with rapamycin were incubated for 12 h, then stained with fluorescent brightener 28 (CFW) and observed by microscopy.

(D) Phenotype of lipid droplets accumulation in the mycelia of the WT strain treated with or without 100 ng/mL rapamycin for 6 h (bar, 10 μm).

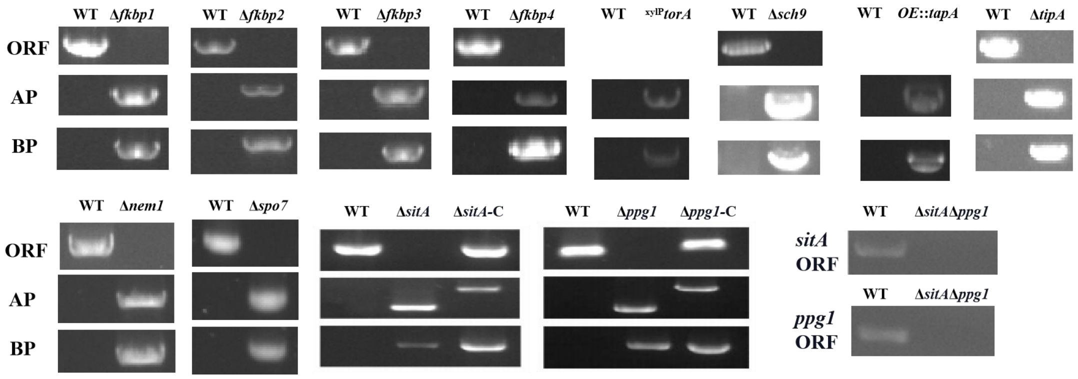


**Figure S2. Construction of all mutants using homologous recombination.** The ORFs of the genes were replaced, which led to the deletion of the genes. All mutations were validated by PCR in *A. flavus*.

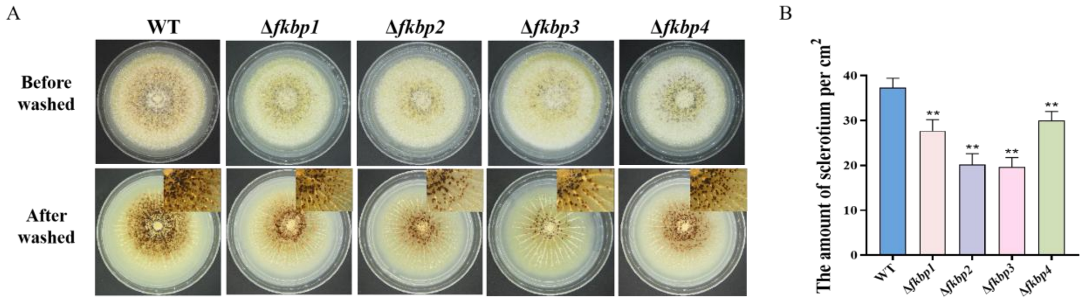


**Figure S3. Fkbps regulate sclerotia biosynthesis in *A. flavus*.**

(A) Phenotypic characterization of WT, Δ*fkbp1*, Δ*fkbp2*, Δ*fkbp3*, and Δ*fkbp4* strains grown on CM medium at 37°C for 7 days.

(B) Quantitative analysis of sclerotium formation as in (A).

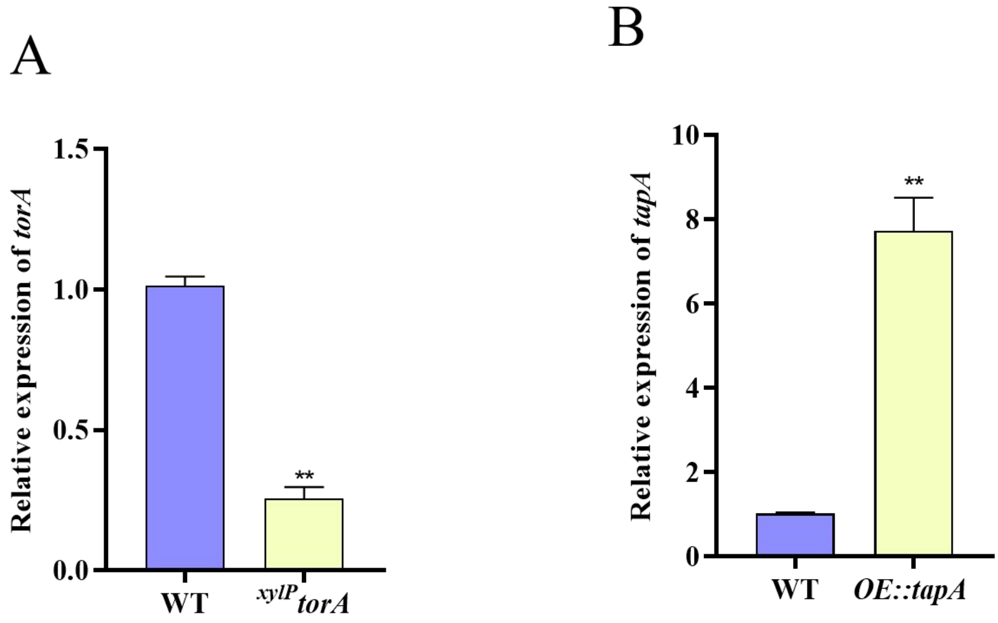


**Figure S4. Transcriptional levels of *tor* and *tapA*.**

(A) Transcriptional levels of *torA* expression in the WT and *$^{xylP}$torA* strains in YXT medium.

(B) Transcriptional levels of the *tapA* gene in the WT and *OE*::*tapA* strains.

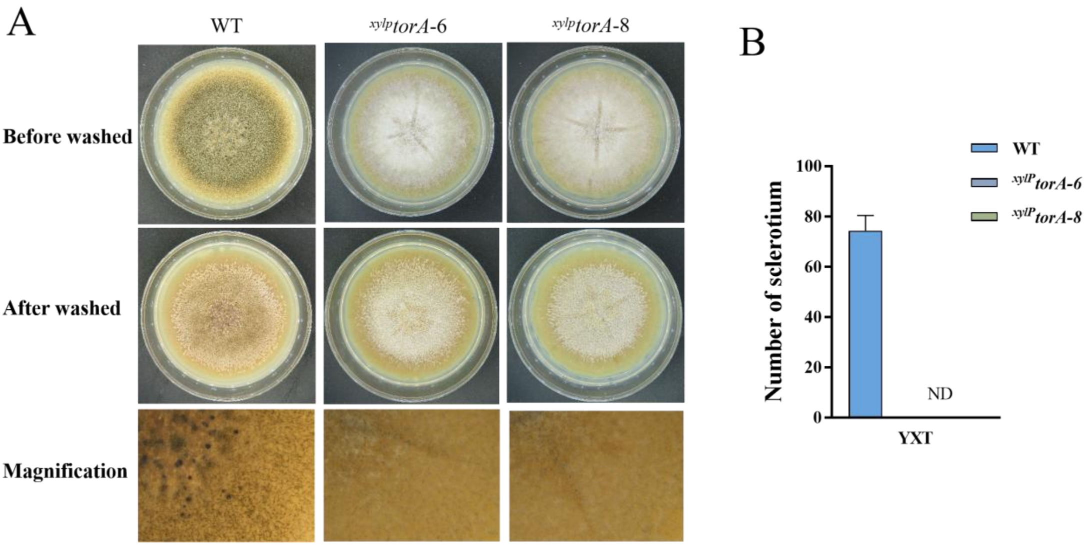


**Figure S5. TorA regulate sclerotia biosynthesis in *A. flavus*.**

(A) Phenotypic characterization of WT and the $^{xylP}$*torA* strains grown on YXTmedium at 37°C for 7 days.

(B) Quantitative analysis of sclerotium formation as in (A).

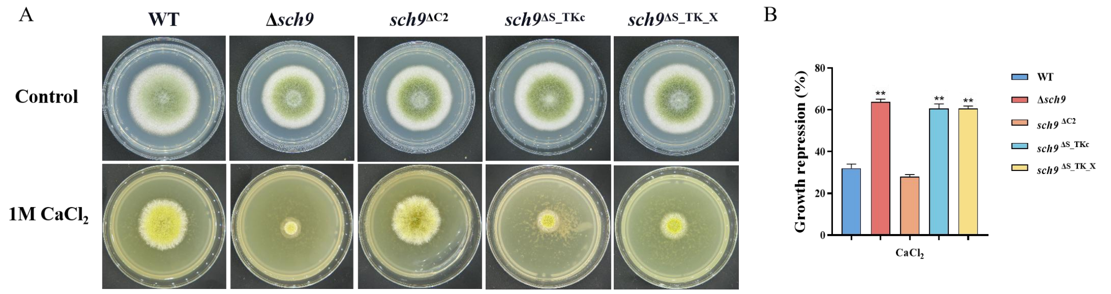


**Figure S6. Sch9 is involved in calcium stress.**

(A) Colony morphology of WT, Δ*sch9*, *Sch9*$^{\Delta C2}$, *Sch9*$^{\Delta S\_TKc}$, and *Sch9*$^{\Delta S\_TK\_X}$ strains on YGT media amended with 1 M $CaCl_2$ for 3 days.

(B) The growth inhibition rate of WT and all mutant strains under calcium stress.

**Table S1**. WT and mutant strains of fungi used in this study.

| Strain | Genotype description | References |
|---|---|---|
| *A. flavus* PTS | Δ*ku70*; Δ*niaD*; Δ*pyrG* | Chang et al, 2010 |
| *A. flavus* wild-type | Δ*ku70*; Δ*niaD*; Δ*pyrG*::*pyrG* | Saved in our lab |
| *A. flavus* Δ*fkbp1* | Δ*ku70*; Δ*niaD*; Δ*fkbp1*::*pyrG* | In study |
| *A. flavus* Δ*fkbp2* | Δ*ku70*; Δ*niaD*; Δ*fkbp2*::*pyrG* | In study |
| *A. flavus* Δ*fkbp3* | Δ*ku70*; Δ*niaD*; Δ*fkbp3*::*pyrG* | In study |
| *A. flavus* Δ*fkbp4* | Δ*ku70*; Δ*niaD*; Δ*fkbp4*::*pyrG* | In study |
| *A. flavus fkbp3* $^{K5A}$ | Δ*ku70*; Δ*niaD*; *fkbp3* $^{K5A}$::*pyrG* | In study |
| *A. flavus fkbp3* $^{K19A}$ | Δ*ku70*; Δ*niaD*; *fkbp3* $^{K19A}$::*pyrG* | In study |
| *A. flavus fkbp3* $^{K40A}$ | Δ*ku70*; Δ*niaD*; *fkbp3* $^{K40A}$::*pyrG* | In study |
| *A. flavus fkbp3* $^{K42A}$ | Δ*ku70*; Δ*niaD*; *fkbp3* $^{K42A}$::*pyrG* | In study |
| *A. flavus fkbp3* $^{K55A}$ | Δ*ku70*; Δ*niaD*; *fkbp3* $^{K55A}$::*pyrG* | In study |
| *A. flavus fkbp3* $^{K65A}$ | Δ*ku70*; Δ*niaD*; *fkbp3* $^{K65A}$::*pyrG* | In study |
| *A. flavus* $^{xylP}$*torA* | Δ*ku70*; Δ*niaD*; *xylP(torA)*::*pyrG* | In study |
| *A. flavus* Δ*sch9* | Δ*ku70*; Δ*niaD*; Δ*sch9*::*pyrG* | In study |
| *A. flavus sch9* $^{\Delta C2}$ | Δ*ku70*; Δ*niaD*; *sch9* $^{\Delta C2}$::*pyrG* | In study |
| *A. flavus sch9* $^{\Delta S\_TKc}$ | Δ*ku70*; Δ*niaD*; *sch9* $^{\Delta S\_TKc}$ ::*pyrG* | In study |
| *A. flavus sch9* $^{\Delta S\_TK\_X}$ | Δ*ku70*; Δ*niaD*; *sch9* $^{\Delta S\_TK\_X}$::*pyrG* | In study |
| *A. flavus sch9* $^{K340A}$ | Δ*ku70*; Δ*niaD*; *sch9* $^{K340A}$::*pyrG* | In study |
| *A. flavus OE::tapA* | *Δku70*; Δ*niaD*; *gpdA(tapA)*:: *pyrG* | In study |

| | | |
|---|---|---|
| *A. flavus* Δ*tipA* | *Δku70*; Δ*niaD*; Δ*tipA*::*pyrG* | In study |
| *A. flavus* Δ*sitA* | *Δku70*; Δ*niaD*; Δ*sitA*::*pyrG* | In study |
| *A. flavus* Δ*ppg1* | *Δku70*; Δ*niaD*; Δ*ppg1*::*pyrG* | In study |
| *A. flavus* Δ*sitA/ ppg1* | *Δku70*; Δ*niaD*; Δ*ppg1*::*pyrG*, Δ*sitA*::*ptr* | In study |
| *A. flavus* Δ*sitA-Com* | Δ*ku70*; Δ*niad*; Δ*sitA*::*pyrG* | In study |
| *A. flavus* Δ*ppg1-Com* | Δ*ku70*; Δ*niad*; Δ*ppg1*::*pyrG* | In study |
| *A. flavus* Δ*nem1* | *Δku70*; Δ*niaD*; Δ*nem1*::*pyrG* | In study |
| *A. flavus* Δ*spo7* | *Δku70*; Δ*niaD*; Δ*spo7*::*pyrG* | In study |

**Table S2**. Primers used in this study.

| Primer | Sequence（5’-3’） |
|---|---|
| Fkbp1 L-F | CAATAGCAGCAACAGCCTCA |
| Fkbp1 L-R | GGGTGAAGAGCATTGTTTGAGGCGCTGGTGGAGTTCGTATCGA |
| Fkbp1 R-F | GCATCAGTGCCTCCTCTCAGACCATCCACTCCTCAATAGACT |
| Fkbp1 R-R | CGAGGCATGATGATATCGAC |
| Fkbp1 C-F | ATGGTGATGGAGTTGAGTTG |
| Fkbp1 C-R | CATTGCGCGCATATCTCAC |
| Fkbp1 O-F | CAAGGTCTCCATCCACTACA |
| Fkbp1 O-R | TGATAATGCGATGCCACTAG |
| Fkbp2 L-F | AGTGGAGTGCAAGGACCTT |
| Fkbp2 L-R | GGGTGAAGAGCATTGTTTGAGGCGCGTGGTTGAGTTAATTGAG |
| Fkbp2 R-F | GCATCAGTGCCTCCTCTCAGACCCACGGTGTTGCTTGGTACT |
| Fkbp2 R-R | TACAGCAAGTACCGAGTGAT |
| Fkbp2 C-F | ATGGTCGGTTCCATGTTCGG |
| Fkbp2 C-R | TAGTCTGCGCATAATGTTGG |
| Fkbp2 O-F | ATGCGTTTCTCAATCTTCTC |
| Fkbp2 O-R | CTCATCCTTCGAAACACCAT |
| FKBP3 L-F | CATCCGTTGTTTATGCTCTG |
| FKBP3 L-R | GGGTGAAGAGCATTGTTTGAGGCGTCGGTACCTGGATTGTGTA |
| FKBP3 R-F | GCATCAGTGCCTCCTCTCAGACTCGAATGCCTAACCTCACCA |
| FKBP3 R-R | TCATTGAAGTCCTTACCTGG |

| | |
|---|---|
| FKBP3 C-F | CATTCAAGAGACCTTCCAGA |
| FKBP3 C-R | GTGAACAGGCACAGGTCCTA |
| FKBP3 O-F | ATGGGTGTCACTAAGACGCT |
| FKBP3 O-R | GCTTTGGCATCTCCTTGTTG |
| FKBP4 L-F | ATCTGATCCACCAGCCTCCA |
| FKBP4 L-R | GGGTGAAGAGCATTGTTTGAGGCATCTGATCCACCAGCCTCCA |
| FKBP4 R-F | GCATCAGTGCCTCCTCTCAGACGTATCACGGTGGCAGTGTT |
| FKBP4 R-R | TCATCACAGCAGTAGGAGTC |
| FKBP4 C-F | CCACGTAGATAGGTGATGCA |
| FKBP4 C-R | AGCTTGAAGGGTCGGAACTA |
| FKBP4 O-F | ATGTCTGTCCAACCTGTCGC |
| FKBP4 O-R | TCACCACCAACGGCCATACC |
| Tor L-F | TGCTATCTCACGAGGTAACAC |
| Tor L-R | GGGTGAAGAGCATTGTTTGAGGCGTGGTCACCATAGGATCGA |
| Tor R-F | GCATCAGTGCCTCCTCTCAGACAGCCCTGGCGGATGATACTA |
| Tor R-R | GTGCTATGAGTTGCGGTGT |
| Tor C-F | GCTTTAGAGTGGCTTCAGTCT |
| Tor C-R | GTCTAAGGTAGCTCTTAGC |
| Tor O-F | AGTTGTACCTGCCACAGCT |
| Tor O-R | ACATTGACTTGCCGTGATAG |
| Sch9 L-F | GACATTCGAATGTTGTCACCGT |
| Sch9 L-R | GGGTGAAGAGCATTGTTTGAGGCTACCAAGCGCAGGTGGAGGGAT |

| | |
|---|---|
| Sch9 R-F | GCATCAGTGCCTCCTCTCAGACTGATCCGATCTAGTATATGC |
| Sch9 R-R | GTTCCTTGATGCGAAGGAAGAG |
| Sch9 C-F | AACCAATAGTAGACCGAAGC |
| Sch9 C-R | CTTGATGGCAAGTGTCAAGG |
| Sch9 O-F | TCTACAGGACGACGACAATC |
| Sch9 O-R | ATAGACCTGACCGAATGTGC |
| TapA L-F | CAAGATTCCTCTATCGGACG |
| TapA L-R | GGGTGAAGAGCATTGTTTGAGGCAGCTGCATGAAACAGAGCTC |
| TapA R-F | GCATCAGTGCCTCCTCTCAGACTCCTCTGGCACATGCTTAAT |
| TapA R-R | TATAAGGTAGTGGAGGCGAC |
| TapA C-F | GAGTTGAGTAGACATTGGTTC |
| TapA C-R | GTCATTTAACCGGTTCTGCT |
| TapA O-F | GCAACGATGTCAATGCCACA |
| TapA O-R | CGTCGATGGTCATAGTAGGC |
| TipA L-F | ACGAGGTCAATGGCGATTCA |
| TipA L-R | GGGTGAAGAGCATTGTTTGAGGCGAATAGATAGCGGAAAGTGC |
| TipA R-F | GCATCAGTGCCTCCTCTCAGACCGCTGTATCAAACTGAGGAG |
| TipA R-R | TATTGTGGCCATAGCAAGGC |
| TipA C-F | CTATGAGGAGTAAGCATGTG |
| TipA C-R | GTCCATGCGATGTTCAGCAT |
| TipA O-F | ATGTCTTCTGAGAACTCTCG |
| TipA O-R | CCTGTCGAACTGTCTCATAG |

| | |
|---|---|
| SitA L-F | CCTTGGTGGCTTTATCCG |
| SitA L-R | GGGTGAAGAGCATTGTTTGAGGCAGGGTCAGAACGGTCA |
| SitA R-F | GCATCAGTGCCTCCTCTCAGACGTTCGGAGAAACCGTTCA |
| SitA R-R | TCCATTAGAACACGAAAGAC |
| SitA C-F | TAAGGGTGTGGTGGTCACGG |
| SitA C-R | GGCCTGGTTCAAGATTACAT |
| SitA O-F | CAAGTATCTTTCAGAGCAGCAT |
| SitA O-R | TTATCTCCAAACAGCCAAC |
| Ppg1 L-F | TAGGCTCCACTCCAAATA |
| Ppg1 L-R | GGGTGAAGAGCATTGTTTGAGGCGTCTGCCAGCAACTCC |
| Ppg1 R-F | GCATCAGTGCCTCCTCTCAGACGTTAGATGATGGGCAAA |
| Ppg1 R-R | GAGGAAGGGTGTTAGAGTTA |
| Ppg1 C-F | TTTCCCGCTCCCTATCCC |
| Ppg1 C-R | CCACCGACAAGCCCGTTT |
| Ppg1 O-F | GGGTTCTGTCCGTCCTTA |
| Ppg1 O-R | CCGTCCATCCACTCCAT |
